# Exercise and disease state influence the beneficial effects of Fn14-depletion on survival and muscle pathology in the *SOD1^G93A^* amyotrophic lateral sclerosis (ALS) mouse model

**DOI:** 10.1101/2024.07.05.602199

**Authors:** Gareth Hazell, Nina Ahlskog, Emma R Sutton, Magnus Okoh, Joseph M Hoolachan, Taylor Scaife, Sara Iqbal, Eve McCallion, Amarjit Bhomra, Anna J Kordala, Frederique Scamps, Cedric Raoul, Matthew JA Wood, Melissa Bowerman

## Abstract

**Background:** Amyotrophic lateral sclerosis (ALS) is a devastating and incurable neurodegenerative disease. Accumulating evidence strongly suggests that intrinsic muscle defects exist and contribute to disease progression, including imbalances in whole-body metabolic homeostasis. We have previously reported that tumour necrosis factor (TNF)-like weak inducer of apoptosis (TWEAK) and fibroblast growth factor inducible 14 (Fn14) are significantly upregulated in skeletal muscle of the *SOD1^G93A^* ALS mouse model. While antagonising TWEAK did not impact survival, we did observe positive effects in skeletal muscle. Given that Fn14 has been proposed as the main effector of the TWEAK/Fn14 activity and that Fn14 can act independently from TWEAK in muscle, we suggest that manipulating Fn14 instead of TWEAK in the *SOD1^G93A^* ALS mice could lead to differential and potentially improved benefits.

**Methods:** We thus investigated the contribution of Fn14 to disease phenotypes in the *SOD1^G93A^* ALS mice. To do so, Fn14 knockout mice (*Fn14^-/-^*) were crossed onto the *SOD1^G93A^* background to generate *SOD1^G93A^;Fn14^-/-^* mice. Investigations were performed on both unexercised and exercised (rotarod and/or grid test) animals (wild type (WT), *Fn14*^-/-^, *SOD1^G93A^* and *SOD1^G93A^;Fn14^-/-^*).

**Results:** Here, we firstly confirm that the TWEAK/Fn14 pathway is dysregulated in skeletal muscle of *SOD1^G93A^* mice. We then show that Fn14-depleted *SOD1^G93A^* mice display an increased lifespan and decreased muscle pathology, without an impact on motor function, and that this is dependent on exposure to exercise. Indeed, we observe that endurance (rotarod) and resistance (grid test) exercises influence the positive effects of Fn14 deletion on survival and muscle phenotypes in *SOD1^G93A^* mice, which may be further influenced by genotype and disease state.

**Conclusions:** Our study provides further insights on the different roles of the TWEAK/Fn14 pathway in pathological skeletal muscle and how they can be influenced by age, disease and metabolic state. This is particularly relevant in the ALS field, where combinatorial therapies that include exercise regimens are currently being explored. As such, a better understanding and consideration of the interactions between treatments, muscle metabolism and exercise will be of importance in future studies.

## BACKGROUND

Amyotrophic lateral sclerosis (ALS) is a devastating and currently incurable neurodegenerative disease. Once symptomatic, the median survival of patients is usually between 3 and 5 years. Clinical manifestations typically occur in mid-life, followed by the rapid and progressive wasting of muscles and subsequent paralysis [1]. ALS can be sporadic (∼80%) or familial (∼20%) [2], and in the latter case can be caused by numerous genetic mutations with the most common being in *chromosome 9 open reading frame 72* (*C9ORF72*) [3,4], *superoxide dismutase 1* (*SOD1*) [5], *Fused in Sarcoma* (*FUS*) [6,7] and *TAR DNA-binding protein 43* (*TDP-43*) [8–10]. Both sporadic and familial ALS patients present similar symptoms and pathophysiology. While the primary pathological target of ALS is undeniably the motor neurons (both upper and lower), accumulating evidence strongly suggests that intrinsic muscle defects exist and contribute to disease progression and presentation [11]. Indeed, the muscle-restricted expression of mutant SOD1 results in a canonical ALS pathophysiology [12,13]. Furthermore, aberrant genetic, biochemical, developmental, regulatory and physiological changes prior to, or accompanying, motor neuron loss are observed in ALS muscle and progenitor cells [11]. As muscle plays an important role in maintaining systemic energy homeostasis [14], intrinsic muscle defects can have severe consequences on whole-body metabolic homeostasis. Interestingly, instances of insulin resistance [15], hyperlipidemia [16], hyperglycemia [17], aberrant fatty acid metabolism [18], hyperglucagonemia [19], glucose intolerance [18] and development of diabetes [20] have all been reported in ALS patients and animal models. Furthermore both dietary and exercise interventions, which are direct modulators of muscle metabolism [21], have been demonstrated to impact disease progression in ALS patients and animal models [22–24]. Thus, uncovering and targeting pathological molecular effectors in ALS muscle may lead to tissue-specific and whole-body improvements [11,25,26].

One important pathway that contributes to skeletal muscle health, function and metabolism is controlled by the binding of the tumour necrosis factor (TNF)-like weak inducer of apoptosis (TWEAK) ligand to the TNF fibroblast growth factor inducible 14 (Fn14) receptor [27,28]. Interestingly, the TWEAK/Fn14 pathway can impact muscle positively or negatively depending on the levels of TWEAK present. High levels are typically associated with detrimental effects while low levels have a beneficial impact [27,28]. Similarly, Fn14 expression is typically very low in healthy muscle and becomes upregulated in muscle atrophy conditions, which can lead to sustained muscle pathology if not restored to normal levels [27,28]. Furthermore, TWEAK and Fn14 have both been implicated in the regulation of key muscle metabolic effectors such as peroxisome proliferative activated receptor, gamma, coactivator 1 alpha (PGC-1α), Slc2a4 solute carrier family 2, member 4 (GLUT-4), hexokinase 2 (HKII) and Krüppel-like transcription factor 15 (KLF15) [29].

What still remains unclear however, is the potential role of the TWEAK/Fn14 pathway in neuromuscular conditions, where chronic muscle wasting occurs due to motor neuron loss and muscle denervation [30]. In an attempt to explore this further, we have previously investigated the TWEAK/Fn14 signalling cascade in mouse models of ALS and spinal muscular atrophy (SMA), a childhood neuromuscular disease [31]. In pre-weaned SMA mice, we observed a significant downregulation of *Tweak* and *Fn14* in various skeletal muscles during disease progression, accompanied by the expected dysregulation of *PGC-1*α, *Glut4*, *HKII* and *Klf15* [32]. Interestingly, administering Fc-TWEAK, an agonist of the pathway, to SMA mice, improved several canonical disease phenotypes [32]. Conversely, we have previously observed that *Tweak* and *Fn14* are significantly upregulated in the skeletal muscle of *SOD1^G93A^* ALS mice during disease progression [33]. While antagonising TWEAK, either genetically or pharmacologically, did not impact survival, we did observe positive effects in skeletal muscle [33]. Since the receptor has been proposed as the main effector of the TWEAK/Fn14 pathway activity [34] and that Fn14 can act independently from TWEAK in muscle [35], it is possible that manipulating Fn14 instead of TWEAK in the *SOD1^G93A^* ALS mice could lead to differential and/or improved benefits.

In this study, we investigated the effect of Fn14 depletion on disease progression and muscle pathology in *SOD1^G93A^* ALS mice by crossing Fn14 knockout mice (*Fn14^-/-^*) with the *SOD1^G93A^*mouse model. We confirmed that the TWEAK/Fn14 pathway is dysregulated in the skeletal muscle of *SOD1^G93A^* mice. We then showed that Fn14-depleted *SOD1^G93A^* mice had an increased lifespan and decreased muscle pathology, which was dependent on exposure to exercise. Our study provides further insights into the different roles of the TWEAK/Fn14 pathway in skeletal muscle and how they may be influenced by age, disease and metabolic state.

## METHODS

### Animals and animal procedures

*SOD1^G93A^* mice (B6.Cg-Tg(SOD1*G93A)1Gur/J) were obtained from Jackson Laboratories (Strain #: 004435). The *Fn14^-/-^*mouse model [36] was provided by Biogen.

Experimental procedures with live animals were authorized and approved by the University of Oxford ethics committee and UK Home Office (Project licenses PDFEDC6F0 and 30/2907) in accordance with the Animals (Scientific Procedures) Act 1986.

For survival studies, mice were weighed and monitored daily and culled upon reaching their defined humane endpoint as specified in the project license.

For all experiments, litters were randomly assigned treatment at birth. Sample sizes were determined based on similar studies with *SOD1^G93A^*mice.

For the grid test, we used our previously described protocol [33], whereby starting with a 40 g metal grid (followed by 30, 20 and 10 g grids), we measured the time (maximum 30 s) the animal held on to the grid before dropping it. The experiment was repeated three times with each grid. Muscle strength (arbitrary units) was quantified with the following formula: (40 g × best time) + (30 g × best time) + (20 g × best time) + (10 g × best time).

For the rotarod test, we followed the previously described protocol [37], whereby mice were placed on the rotarod (opposite orientation to rotation) with an acceleration protocol of 4 to 40 rpm in 300 s. The latency to fall (s) and highest rpm reached was recorded.

To reduce the total number of mice used, the fast-twitch *tibialis anterior* (TA) and gastrocnemius muscles from the same mice were used for molecular and histological analyses, respectively.

### qPCRs

RNA was extracted from tissues with the RNeasy kit (Qiagen) or with a Isolate II RNA Mini Kit (Bioline) as per the manufacturers’ instructions. The same RNA extraction method was employed for similar experiments and equal RNA amounts were used between samples within the same experiments. cDNA was prepared with the High-capacity cDNA Kit (Life Technologies) or qPCRBIO cDNA Synthesis Kit (PBCR Biosystems) according to the manufacturers’ instructions. The same reverse transcription method was employed for similar experiments. The cDNA template was amplified on a StepOnePlus Real-Time PCR Thermocycler (Life Technologies) with SYBR Green Mastermix (Applied Biosystems) or with qPCRBIO SyGreen Blue Mix Hi-ROX (PCR Biosystems). The same amplification method was used for similar experiments. qPCR data was analysed using the StepOne Software v2.3 (Applied Biosystems). Primers used for qPCR were obtained from IDT and sequences for primers were self-designed (Supplementary Table 1). Relative gene expression was quantified using the Pfaffl method [38] and primer efficiencies were calculated with the LinRegPCR software. The relative expression of all genes of interest was normalised to the expression of *RNA polymerase II polypeptide J* (*PolJ*) [39].

### Immunoblots

Freshly prepared RIPA buffer (50 mM Tris pH 8.8, 150mM NaCl, 1% NP-40, 0.5% sodium deoxycholate, 0.1% sodium dodecyl-sulphate (SDS) and complete mini-proteinase inhibitors (Roche)) was used to homogenize tissue. Equal amounts of total protein were loaded in the wells, as measured by Bradford Assay. Protein samples were first diluted 1:1 with Laemmli sample buffer (Bio-Rad, Hemel Hempstead, UK) containing 5% β-mercaptoethanol (Sigma) and heated at 100°C for 10 minutes. Next, samples were loaded on freshly made 1.5 mm 12% polyacrylamide separating and 5% stacking gel and electrophoresis was performed at 120 V for ∼1.5 hours in running buffer. Proteins were then transferred from the gel onto to a polyvinylidene fluoride membrane (Merck Millipore) via electroblotting at 120 V for 60 minutes in transfer buffer containing 20% methanol. Membranes were then incubated for 2 hours in Odyssey Blocking Buffer (Licor). The membrane was then probed overnight at 4°C with the primary antibodies (p105/p50, Abcam ab32360, 1:1000; Actin, Abcam ab3280, 1:1000) in Odyssey Blocking Buffer and 0.1% Tween-20. The next day, after three 10-minute washes in phosphate buffered saline (PBS), the membrane was incubated for 1 hour at room temperature with secondary antibodies (goat anti-rabbit IgG 680RD, LI-COR 926-68071, 1:1000,; goat anti-mouse IgG 800CW, LI-COR, 926-32210, 1:1000). Lastly, the membrane was washed three times for 10 minutes in PBS and visualized by scanning the 700 nm and 800 nm channels on the LI-COR Odyssey CLx infrared imaging system (LI-COR) for 2.5 minutes per channel. The background was subtracted and signal of protein of interest was divided by signal of the housekeeping protein (actin).

### Laminin staining of skeletal muscles

*Tibialis anterior* (TA) muscles were fixed in 4% paraformaldehyde (PFA) overnight. Tissues were sectioned (13 μm) and incubated in blocking buffer (0.3% Triton-X, 20% foetal bovine serum (FBS) and 20% normal goat serum in PBS) for 2 hours. After blocking, tissues were stained overnight at 4°C with rat anti-laminin (Sigma L0663, 1:1000) in blocking buffer. The next day, tissues were washed in PBS and probed using a goat-anti-rat IgG 488 secondary antibody (Invitrogen A-11006, 1:500) for one hour. PBS-washed tissues were mounted in Fluoromount-G (Southern Biotech). Images were taken with a DM IRB microscope (Leica) with a 20x objective. Quantitative assays were performed blinded on 3-5 mice for each group and five sections per mouse. Myofiber area was measured using Fiji (ImageJ) [40].

### Endplate staining of skeletal muscles

Endplates were stained as previously described [41]. Briefly, whole TA muscle was harvested and fixed in 4% PFA for 15 min. Muscles were incubated with α-bungarotoxin (α-BTX) conjugated to tetramethylrhodamine (BT00012, Biotium, 1:100) at RT for 30Lminutes with ensuing PBS washes. Finally, 2–3 thin filets per muscle were sliced and mounted in Fluoromount-G (Southern Biotech). Images were taken with a confocal microscope, with a 20X objective. The experimenter quantifying endplate size was blinded to the genotype of the animals until all measurements were finalized.

### Statistical Analyses

All statistical analyses were done with the most up to date GraphPad Prism software at time of writing. When appropriate, a Student’s unpaired two-tail *t*-test, a one-way analysis of variance (ANOVA) or a two-way ANOVA was used. Post-hoc analyses used are specified in Figure Legends. Outliers were identified via the Grubbs’ test. For the Kaplan-Meier survival analysis, the log-rank test was used and survival curves were considered significantly different at *p* < 0.05.

## RESULTS

### The Fn14 signalling cascade is dysregulated in skeletal muscle of SOD1^G93A^ mice during disease progression

We have previously demonstrated that *Fn14* mRNA levels significantly increase in the skeletal muscle of *SOD1^G93A^* mice during disease progression, while *Tweak* mRNA levels remained relatively unchanged [33]. Furthermore, genetic and pharmacological reduction of TWEAK activity improved muscle pathology in *SOD1^G93A^*mice [33]. Comparison of Fn14 expression in the skeletal muscle of 20-week-old symptomatic *SOD1^G93A^* and *SOD1^G93A^;Tweak^-/-^*mice showed no significant difference in *Fn14* mRNA expression (Figure 1A). This suggests that genetically depleting the ligand (TWEAK) was not sufficient to reduce the expression of the receptor (Fn14). Since Fn14 is a key factor in modulating the activity of the TWEAK/Fn14 pathway [34], its persistent expression despite Tweak depletion may have limited the benefits on muscle pathology and disease progression.

**Figure 1.**
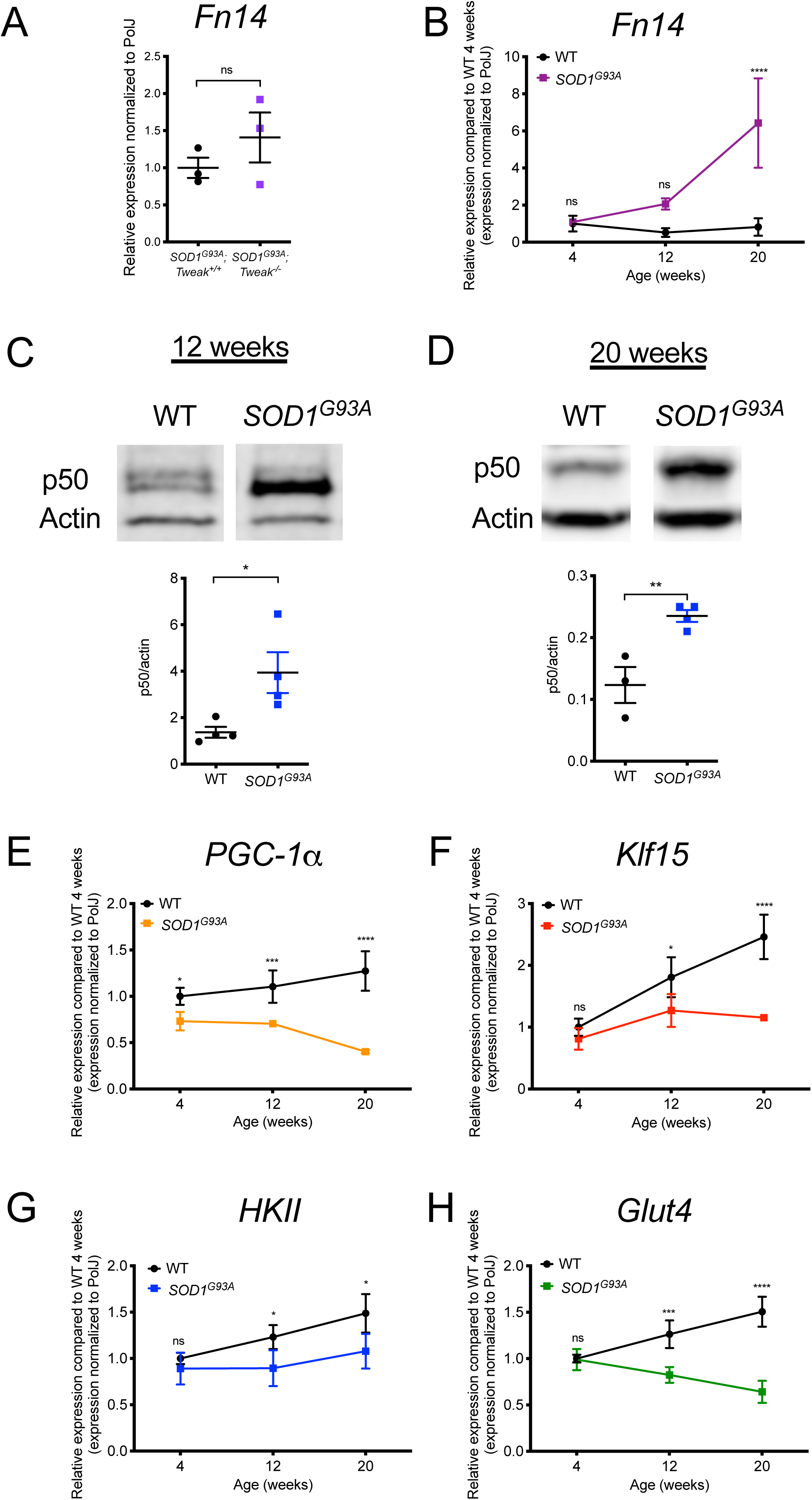
Aberrant expression of the TWEAK/Fn14 signaling pathway Fn14 in skeletal muscle of *SOD1^G93A^*ALS mice. **A)** qPCR analysis of *Fn14* mRNA expression in gastrocnemius muscle from 20-week-old *SOD1^G93A^;Tweak^+/+^*and *SOD1^G93A^;Tweak^-/-^* males. Normalized relative expressions are compared to *SOD1^G93A^;Tweak^+/+^*. Data are scatter dot plot mean ± SEM, *n* = 3 animals per genotype, unpaired *t test*; ns, not significant. **B)** qPCR analysis of *Fn14* mRNA expression in the *tibialis anterior* (TA) of *SOD1^G93A^* and wild type (WT) mice at 4 (pre-symptomatic), 12 (early symptomatic) and 20 (late symptomatic) weeks. Normalized relative expressions are compared to WT 4 weeks. Data are mean ± SEM, *n* = 3-4 animals per experimental group, two-way ANOVA, \*\*\*\**p* < 0.0001. **C-D)** Quantification of NF-κB p50/actin protein levels in the TAs of 12-(**C**) and 20-week-old (**D**) *SOD1^G93A^*and WT. Images are representative immunoblots. Data are scatter dot plot mean ± SEM, *n* = 3-4 animals per experimental group, unpaired *t test*, *p* = 0.0302 (12 weeks), *p* = 0.0088 (20 weeks). **E-F)** qPCR analysis of *PGC-1*α (**E**), *Klf15* (**F**), *HKII* (**G**) and *Glut4* (**H**) mRNAs in TAs of 4-, 12- and 20- week-old *SOD1^G93A^*and WT. Normalized relative expressions are compared to WT 4 weeks. Data are mean ± SEM, n = 3-4 animals per experimental group, two-way ANOVA, \**p* < 0.05, \*\*\**p* < 0.001.

We thus set out to further characterize the Fn14 signalling cascade in skeletal muscle of *SOD1^G93A^* males. We started by reproducing our previously published data [33] and demonstrated that *Fn14* mRNA levels in the *tibialis anterior* (TA) of *SOD1^G93A^* and wild type (WT) mice are similar in 4- (pre-symptomatic) and 12-week-old (early symptomatic) animals while there is a significant increase in 20-week-old (late symptomatic) *SOD1^G93A^* mice (Figure 1B). We next assessed the expression of nuclear factor kappa-light-chain-enhancer of activated B cells (NF-κB) subunit p50, a direct downstream effector of TWEAK/Fn14 signalling in skeletal muscle [28,42] that mediates pathological events in muscle when chronically activated [43]. We found that the expression of NF-κB subunit p50 was significantly upregulated in the TAs of *SOD1^G93A^* mice at both early symptomatic (Figure 1C) and late symptomatic (Figure 1D) time-points compared to WT animals, supporting an increased activity of TWEAK/Fn14 activity in skeletal muscle of ALS mice. Next, we evaluated the gene expression of peroxisome proliferative activated receptor, gamma, coactivator 1 alpha (*PGC-1*α), Krüppel-like transcription factor 15 (*Klf15*), hexokinase 2 (*HKII*) and Slc2a4 solute carrier family 2, member 4 (*Glut4*), metabolic effectors whose expression was previously shown to be inversely correlated to TWEAK/Fn14 activity [29]. Interestingly, we observed a significant decrease in the expression of *PGC-1*α (Figure 1E), *Klf15* (Figure 1F), *HKII* (Figure 1G) and *Glut4* (Figure 1H) in the TA muscles of 12- and 20-week-old *SOD1^G93A^*mice compared to WT animals, providing further support for increased Fn14 expression in *SOD1^G93A^* mice.

Together, our results demonstrate an aberrant hyperactivity of TWEAK/Fn14 signalling in the skeletal muscle of *SOD1^G93A^* mice, impacting key regulatory downstream effectors known to influence overall skeletal muscle health and metabolic homeostasis.

### Genetic deletion of Fn14 increases survival of SOD1^G93A^ mice

We sought to determine if decreasing TWEAK/Fn14 activity in *SOD1^G93A^*mice would improve muscle health and slow disease progression. As described above, we have previously modulated TWEAK activity both genetically and pharmacologically [33]. We thus decided to investigate the impact of depleting the activity of the receptor to abolish downstream signalling effector of the TWEAK/Fn14 pathway [34]. We crossed *SOD1^G93A^* mice with *Fn14^-/-^* mice [36], to generate ALS mice with a homozygous deletion of Fn14. Interestingly, we found that *SOD1^G93A^;Fn14^-/-^*mice had a significantly increased lifespan compared to *SOD1^G93A^* mice (females and males combined) (Figure 2A) without any substantial improvements in weight (Figure 2B,C). In fact, *SOD1^G93A^;Fn14^-/-^*females tended to weigh less than *SOD1^G93A^* females (Figure 2B), while there were no significant differences between *SOD1^G93A^;Fn14^-/-^* and *SOD1^G93A^* males (Figure 2C). Nevertheless, Fn14 depletion appears to have an overall positive impact on disease progression in *SOD1^G93A^*mice.

**Figure 2.**
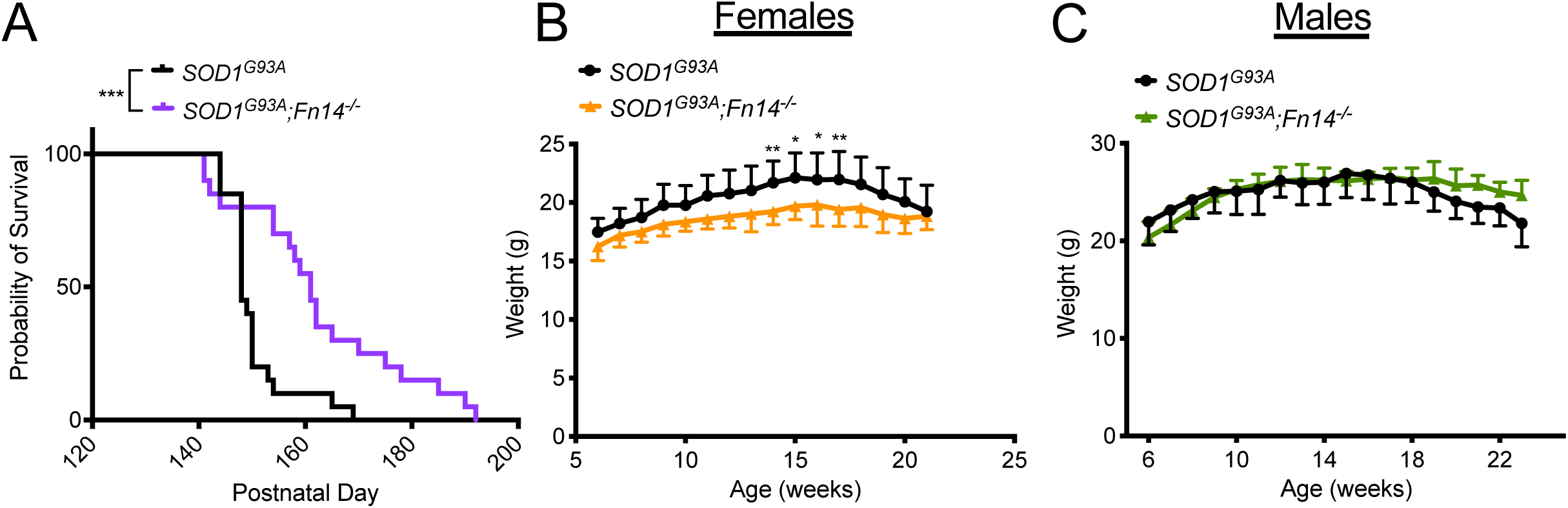
Genetic deletion of *Fn14* increases survival of *SOD1^G93A^* ALS mice. A) Survival curves of untreated *SOD1^G93A^* and *SOD1^G93A^;Fn14^-/-^*mice (males and females combined). Data are represented as Kaplan-Meier survival curves, *n* = 20 animals per experimental group, Log-rank (Mantel-Cox), *p* = 0009. **B-C)** Weekly weights of *SOD1^G93A^* and *SOD1^G93A^;Fn14^-/-^* females (**B**) and males (**C**) from 6 weeks to humane endpoint. Data are meanL± SEM, *n* = 9-11 animals per experimental group, two-way ANOVA, \**p* < 0.05, \*\**p* < 0.01.

### Genetic deletion of Fn14 improves muscle pathology in SOD1^G93A^ mice

We next determined the impact of Fn14 depletion on previously characterised skeletal muscle pathologies in 20-week-old *SOD1^G93A^*males. We firstly measured the myofiber area in the gastrocnemius muscle of WT, *Fn14^-/-^*, *SOD1^G93A^* and *SOD1^G93A^;Fn14^-/-^*mice as muscle wasting is evident in these ALS mice at that symptomatic time-point [33]. We found that while Fn14 depletion in WT animals had no impact on myofiber size, there was a significant increase in myofiber size in *SOD1^G93A^;Fn14^-/-^* mice compared to *SOD1^G93A^* animals (Figure 3A-C).

**Figure 3.**
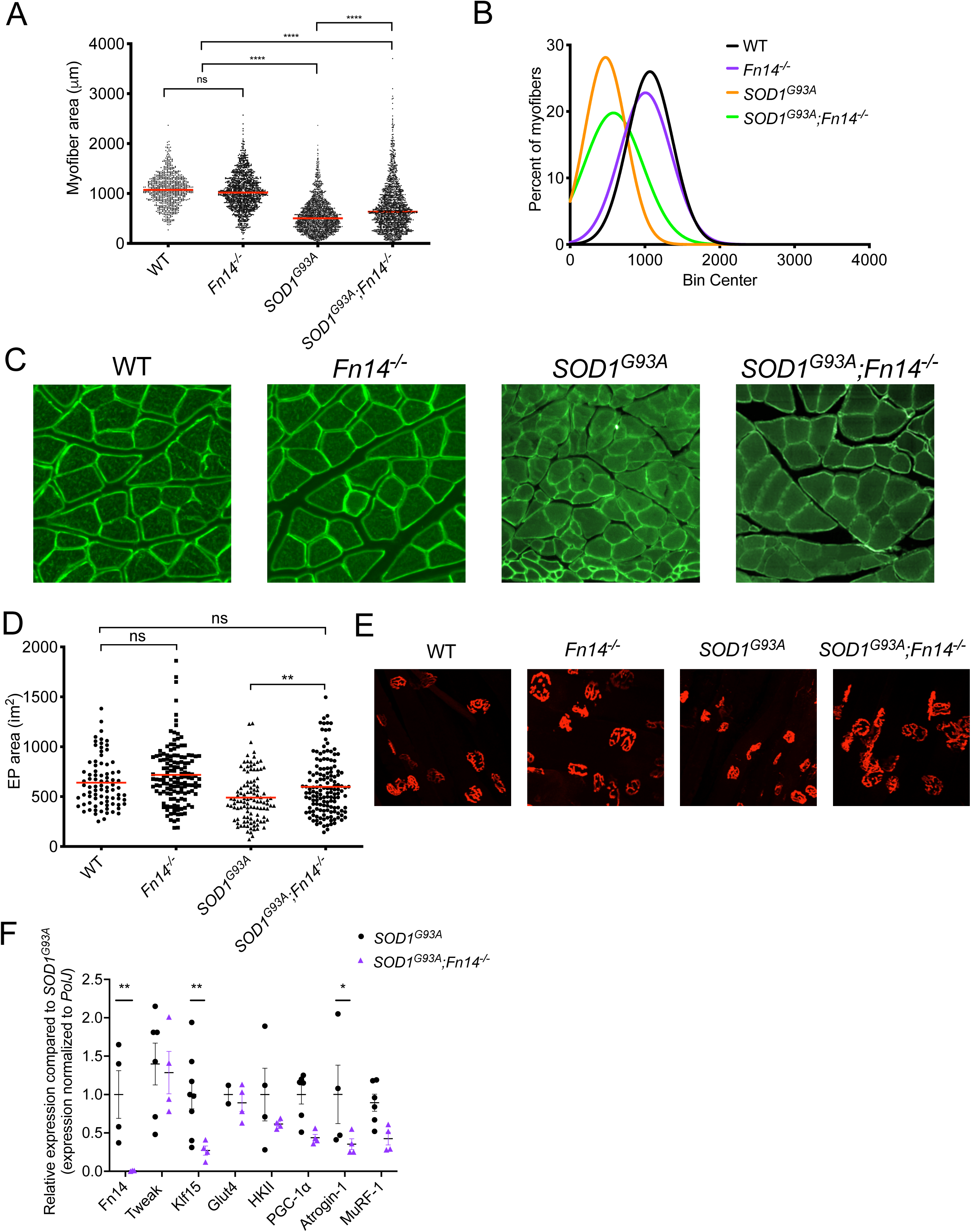
Genetic deletion of *Fn14* improves muscle phenotypes in *SOD1^G93A^* ALS mice. **A)** Quantification of myofiber area of laminin-stained cross-sections of gastrocnemius muscles from 20-week-old WT, *Fn14^-/-^*, *SOD1^G93A^*and *SOD1^G93A^;Fn14^-/-^* males. Data are dot plot and mean, *n* = 3-4 animals per experimental group (>800 myofibers per experimental group), one-way ANOVA, ns = not significant, \*\*\*\**p* < 0.0001. **B)** Relative frequency distribution of myofiber size in gastrocnemius muscles from 20-week-old WT, *Fn14^-/-^*, *SOD1^G93A^*and *SOD1^G93A^;Fn14^-/-^* mice. **C)** Representative images of laminin-stained cross-sections of gastrocnemius muscles from 20-week-old WT, *Fn14^-/-^*, *SOD1^G93A^* and *SOD1^G93A^;Fn14^-/-^* mice. **D)** Quantification of neuromuscular junction endplate (EP) area of alpha-bungarotoxin-stained TA muscles from 20-week-old WT, *Fn14^-/-^*, *SOD1^G93A^* and *SOD1^G93A^;Fn14^-/-^* mice. Data are dot plot and mean, *n* = 3-4 animals per experimental group (>80 myofibers per experimental group), one-way ANOVA, ns = not significant, \*\**p* < 0.01. **E)** Representative images of alpha-bungarotoxin-stained TA muscles from 20-week-old WT, *Fn14^-/-^*, *SOD1^G93A^* and *SOD1^G93A^;Fn14^-/-^* mice. **F)** qPCR analysis of *Fn14*, *Tweak*, *Klf15*, *Glut4*, *HKII*, *PGC-1*α, *Atrogin-1* and *MuRF-1* mRNA in TA muscles from *SOD1^G93A^* and *SOD1^G93A^;Fn14^-/-^*mice. Normalized relative expressions are compared to *SOD1^G93A^*for each gene. Data are scatter dot plot mean L± SEM, n = 3-8 animals per experimental group, two-way ANOVA, \**p* < 0.05.

We also investigated the impact of Fn14 deletion on post-synaptic neuromuscular junction (NMJ) pathologies by evaluating endplate size, which is typically reduced in ALS mice [44] and associated with muscle size [45]. Similar to myofiber size, we observed that Fn14 depletion did not influence the NMJ endplate size in the TA muscles of WT animals, but it significantly increased endplate size in *SOD1^G93A^* mice (Figure 3D-E).

To determine the impact of Fn14 deletion at a molecular level, we investigated the gene expression of molecular effectors associated with the TWEAK/Fn14 signalling cascade (*Fn14*, *Tweak*, *Klf15*, *Glut4*, *HKII* and *PGC-1*α) [29] and muscle atrophy markers (*Atrogin-1* and *MuRF-1*) [46]. We found that the complete elimination of Fn14 in TA muscles of *SOD1^G93A^* mice did not influence the expression of *Tweak*, *Glut4*, *HKII*, *PGC-1*α and *MuRF-1* (Figure 3F). However, we observed a significant decrease in the expression of *Klf15* and, importantly, the atrogene *Atrogin-1* (Figure 3F).

Combined, our analyses of symptomatic mice reveal that deletion of *Fn14* in *SOD1^G93A^* mice improves several muscle wasting phenotypes. This suggests that the aberrant increased expression of Fn14 in skeletal muscle of *SOD1^G93A^* animals may contribute to the muscle pathologies that the disease.

### Enhanced physical activity and Fn14 depletion both have positive effects on survival of SOD1^G93A^ mice

We assessed if the observed molecular and histological benefits in the muscles of Fn14-depleted ALS mice translated into improved motor performance. *SOD1^G93A^* and *SOD1^G93A^;Fn14^-/-^*mice therefore performed a weekly rotarod [47] and grid test [33,48], starting at 8 weeks and ending when the animals reached their defined humane endpoint. Both tests have previously been used in *SOD1^G93A^* mice [33,49] and are aimed at evaluating motor balance and coordination (rotarod) and strength (grid test). We found that there was no significant difference in the time spent on the rotarod between *SOD1^G93A^* and *SOD1^G93A^;Fn14^-/-^*females and males mice (Figure 4A-B). With the grid test, no significant difference in muscle strength was observed between *SOD1^G93A^* and *SOD1^G93A^;Fn14^-/-^*females (Figure 4C), while *SOD1^G93A^* males were significantly stronger than *SOD1^G93A^;Fn14^-/-^*males at the very early pre-symptomatic time-points (Figure 4D). Although these results suggest that Fn14 depletion does not enhance muscle strength and/or performance in *SOD1^G93A^* mice, this might be due to the independent benefits provided by the weekly rotarod and grid tests exercises. Indeed, exercised *SOD1^G93A^* animals had a significantly greater lifespan than unexercised *SOD1^G93A^* mice (Figure 4E). As such, exercised *SOD1^G93A^* and *SOD1^G93A^;Fn14^-/-^*mice had similar survivals, suggesting that both exercise and Fn14 depletion can improve survival in *SOD1^G93A^* mice. Of note, while the median lifespan of exercised *SOD1^G93A^* and *SOD1^G93A^;Fn14^-/-^*mice were not significantly different, there did appear to be a delay in the early deaths in the exercised *SOD1^G93A^;Fn14^-/-^* group, pointing towards a potential combination of independent and dependent mechanisms.

**Figure 4.**
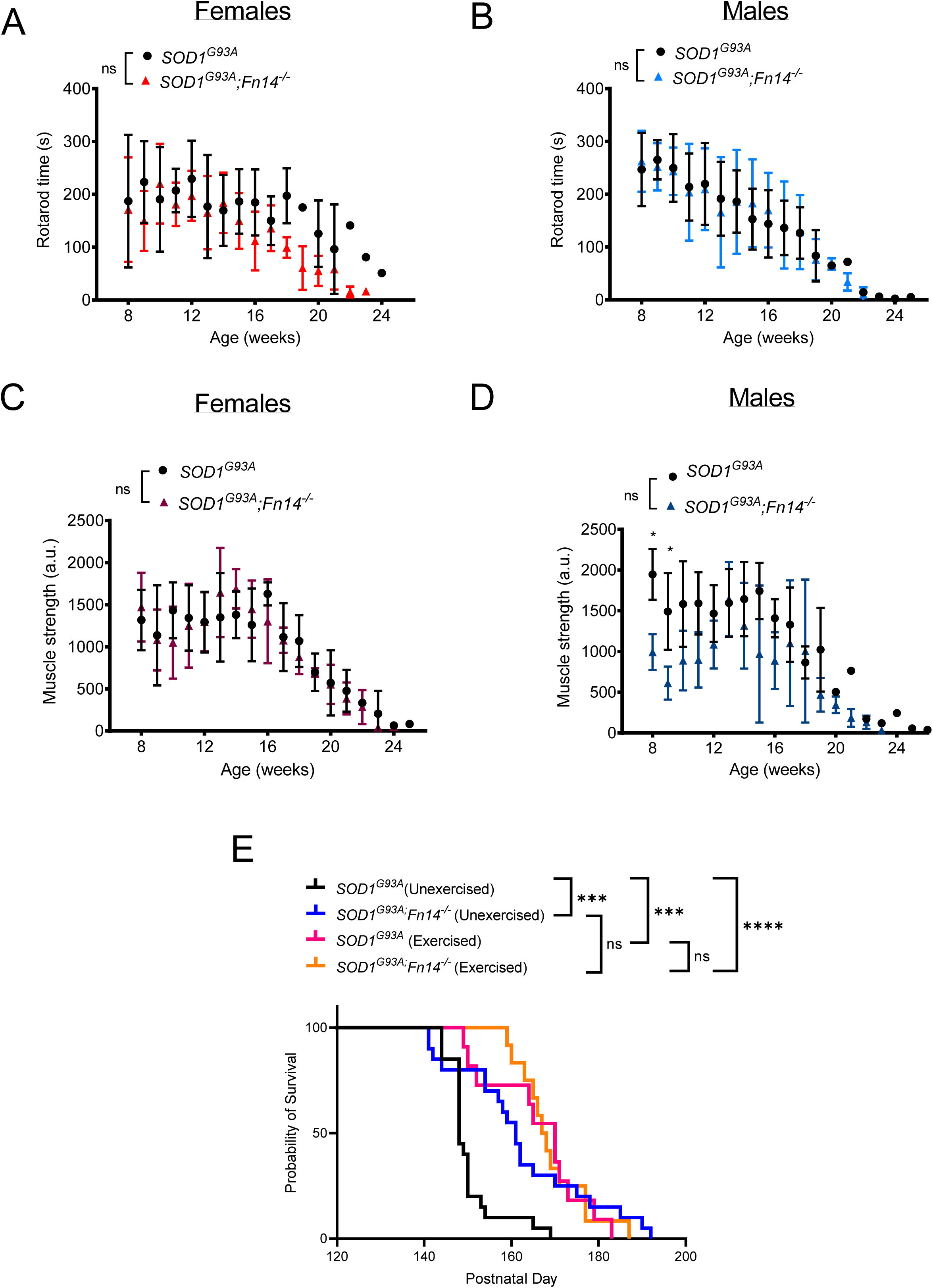
Weekly exercise tests do not reveal any improvements in motor function of Fn14-depleted *SOD1^G93A^*mice and induce benefits on lifespan independent of Fn14 depletion. *SOD1^G93A^*and *SOD1^G93A^;Fn14^-/-^* mice performed both the rotarod and grid test weekly from 8 weeks to humane endpoint. **A-B**) Time in seconds (s) spent on rotarod before falling (maximum 300 s) for *SOD1^G93A^* and *SOD1^G93A^;Fn14^-/-^* female (**A**) and male (**B**) mice. Data are meanL± SEM, *n* = 5-7 animals per experimental group, two-way ANOVA, ns = not significant. **C-D**) Muscle strength (arbitrary units (a.u.) for *SOD1^G93A^* and *SOD1^G93A^;Fn14^-/-^*female (**C**) and male (**D**) mice. Data are meanL± SEM, *n* = 5-7 animals per experimental group, two-way ANOVA, ns = not significant, \**p* < 0.05. **E)** Survival curves of *SOD1^G93A^* and *SOD1^G93A^;Fn14^-/-^*mice that performed both the rotarod and grid test weekly from 8 weeks to humane endpoint (males and females combined). Data are represented as Kaplan-Meier survival curves, *n* = 11-12 animals per experimental group, Log-rank (Mantel-Cox), ns = not significant, \*\*\**p* < 0.001, \*\*\*\**p* < 0.0001.

### Fn14 depletion changes molecular response of SOD1^G93A^ muscle to exercise

To further elucidate the potential complex interactions between exercise, disease state and Fn14 depletion, 12-week-old male mice underwent the rotarod (endurance exercise) or grid test (resistance exercise) for 5 consecutive days. The 12-week time point was chosen as it is an early symptomatic age for *SOD1^G93A^*mice that still allows them to complete both exercise regimens to the same extent as WT and *Fn14^-/-^* animals. The TAs were harvested 2 hours after the last test and compared to those of unexercised sex- and age-matched mice for the expression of the TWEAK/Fn14 metabolic effectors and atrogenes investigated above.

First, we assessed and compared TAs from unexercised and rotarod-trained mice. Interestingly, we observed that *Fn14* expression was significantly upregulated in rotarod-trained *SOD1^G93A^* mice compared to unexercised *SOD1^G93A^*animals, while *Fn14* levels remained unchanged in WT animals (Figure 5A), suggesting a yet to be determined role for Fn14 in exercised *SOD1^G93A^* muscle. Next, we compared the expression of *Tweak*, *MuRF-1*, *Atrogin-1*, *Glut4*, *Klf15*, *HKII* and *PGC-1*α in unexercised and rotarod-trained WT, *Fn14^-/-^*, *SOD1^G93A^*and *SOD1^G93A^;Fn14^-/-^* mice. We found that *Tweak* was significantly increased only in rotarod-trained *SOD1^G93A^;Fn14^-/-^*animals compared to unexercised mice (Figure 5B), suggesting a compensatory mechanism that is plausibly due to reduced levels of its ligand and exercise. The atrogene *MuRF-1* was significantly increased only in the muscles of rotarod-trained *SOD1^G93A^*mice compared to unexercised animals (Figure 5C), indicating that depletion of Fn14 prevents exercise-induced *MuRF-1* upregulation. However, this effect appears to be specific to *MuRF-1* as *Atrogin-1*, which is significantly upregulated in rotarod-trained *SOD1^G93A^* mice, was also increased in rotarod-trained *Fn14^-/-^* and *SOD1^G93A^;Fn14^-/-^*mice compared to unexercised cohorts (Figure 5D). Similarly, the expression of *Glut4* was significantly increased only in rotarod-trained *SOD1^G93A^*mice compared to unexercised animals and remained unchanged in Fn14-depleted groups (Figure 5E). Interestingly, the expression of *Klf15* was significantly upregulated only in rotarod-trained *Fn14^-/-^* animals compared to unexercised mice (Figure 5F). As for the expression of *HKII*, it was significantly increased in rotarod-trained *SOD1^G93A^*and *Fn14^-/-^* mice, while it remained unchanged in rotarod-trained WT mice and *SOD1^G93A^;Fn14^-/-^* compared to unexercised groups (Figure 5G). Finally, the expression of *PGC-1*α was significantly upregulated only in rotarod-trained *SOD1^G93A^*and *SOD1^G93A^;Fn14^-/-^* animals compared to unexercised mice (Figure 5H).

**Figure 5.**
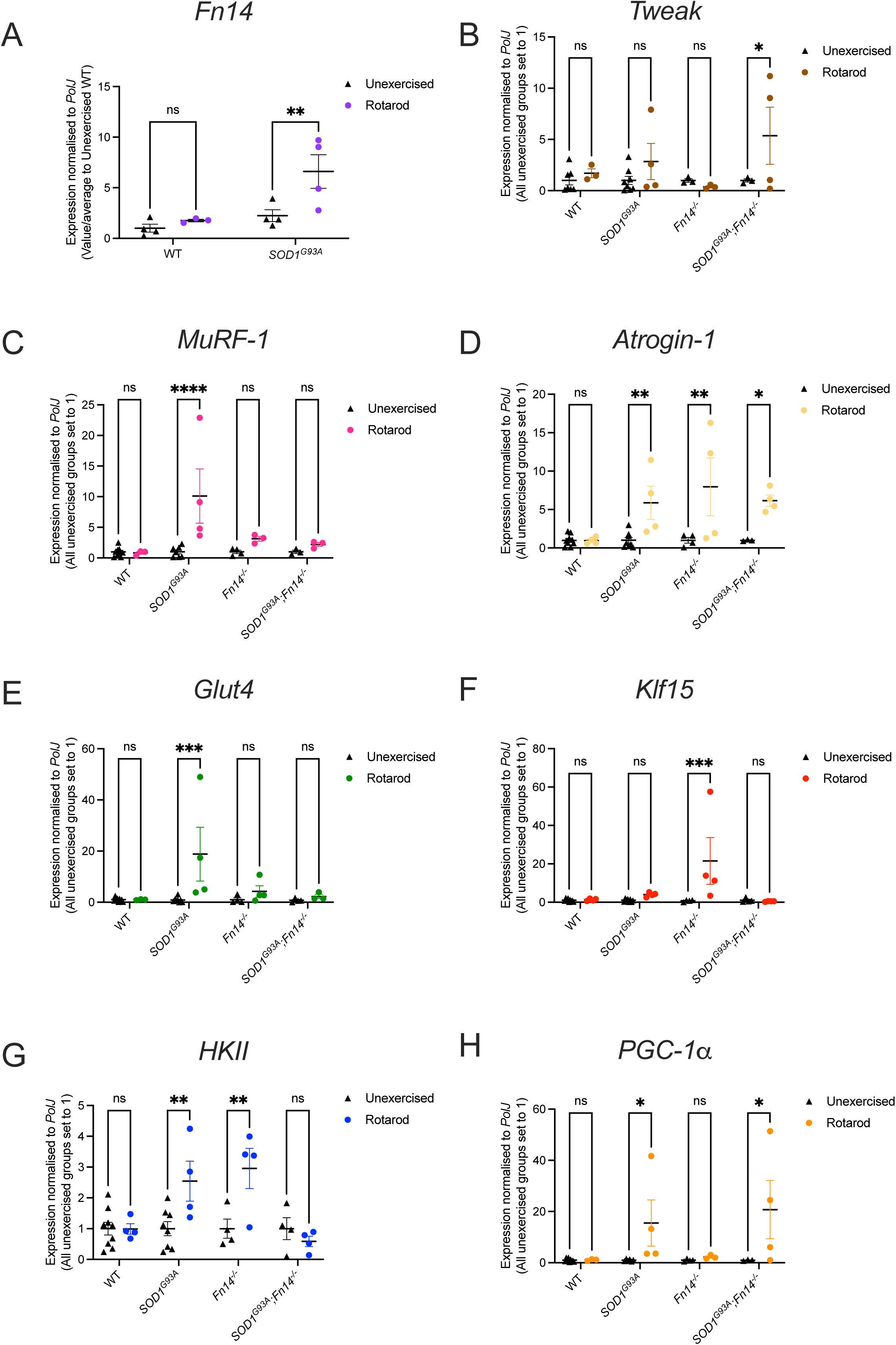
Both rotarod exercise and genotype impact the expression of *Tweak*, *Fn14* and their downstream effectors. 12-week-old WT, *Fn14^-/-^*, *SOD1^G93A^*and *SOD1^G93A^;Fn14^-/-^* males were placed on the rotarod daily for 5 consecutives days. *Tibialis anterior* (TA) muscles were harvested approximately 2 hours after the last bout of exercise. **A**) qPCR analysis of *Fn14* mRNA expression in unexercised and rotarod-exercised WT and *SOD1^G93A^* mice. Data are scatter dot plot mean L± SEM, *n* = 3-4 animals per experimental group, two-way ANOVA, ns = not significant, \*\**p* < 0.01. **B-H**) qPCR analysis of *Tweak* (**B**), *MuRF-1* (**C**), *Atrogin-1* (**D**), *Glut4* (**E**), *Klf15* (**F**), *HKII* (**G**), *PGC-1*α (**H**) mRNA expression in unexercised and rotarod-exercised WT, *Fn14^-/-^*, *SOD1^G93A^* and *SOD1^G93A^;Fn14^-/-^* mice. Data are scatter dot plot mean L± SEM, *n* = 4-9 animals per experimental group, two-way ANOVA, ns = not significant, \**p* < 0.05, \*\*\**p* < 0.001, \*\*\*\**p* < 0.0001.

Next, we performed the same investigations in TAs from unexercised and grid test-trained males. Contrary to what was observed in rotarod-trained *SOD1^G93A^* males (Figure 5A), we found that grid test-trained *SOD1^G93A^* mice expressed significantly less *Fn14* than unexercised *SOD1^G93A^* animals (Figure 6A), suggesting a distinct response between endurance (rotarod) and resistance (grid test) types of activities. On the other hand, *Tweak* expression was significantly increased only in grid test-trained WT animals compared to unexercised mice, while it remained unchanged in grid test-trained animals of the same genotype (Figure 6B). The expression of both atrogenes, *MuRF-1* and *Atrogin-1*, was significantly upregulated only in grid-test trained *SOD1^G93A^* mice compared to unexercised animals and restored to low levels when *Fn14* was depleted (Figure 6C-D). *Glut4* levels were unchanged in all experimental groups when comparing unexercised animals to grid test-trained mice (Figure 6E). Interestingly, *Klf15* expression was significantly upregulated only in *SOD1^G93A^;Fn14^-/-^* animals compared to unexercised mice, as it remained unchanged in all other groups (Figure 6F). Similar to *Glut4*, *HKII* levels were also unchanged in all experimental groups when comparing unexercised animals to grid test-trained mice (Figure 6G). Finally, *PGC-1*α expression was significantly increased only in grid test-trained *SOD1^G93A^* mice compared to unexercised animals and returned to lower levels in Fn14-depleted animals (Figure 6H).

**Figure 6.**
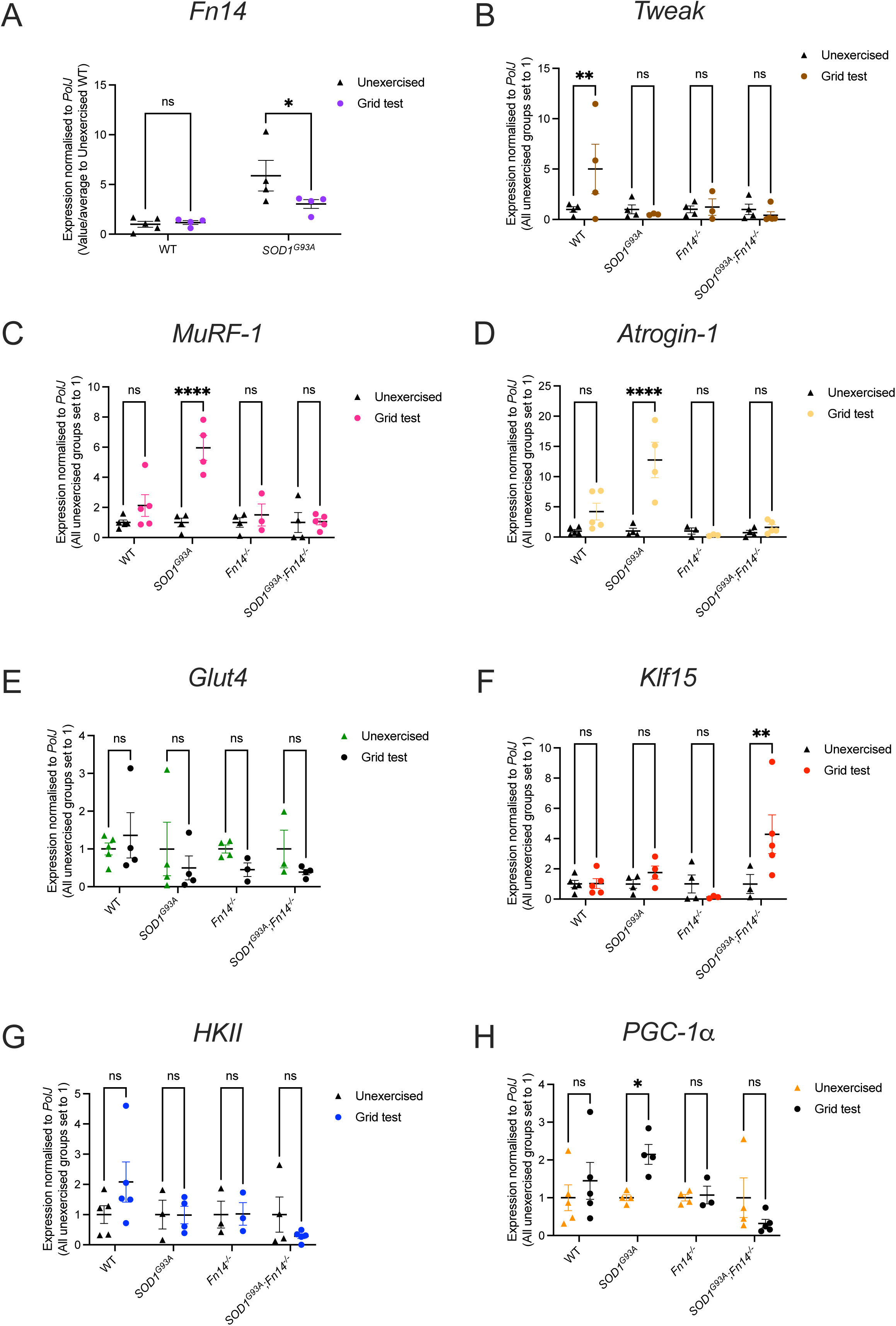
Both grid test exercise and genotype impact the expression of *Tweak*, *Fn14* and their downstream effectors. 12-week-old WT, *Fn14^-/-^*, *SOD1^G93A^* and *SOD1^G93A^;Fn14^-/-^* males performed the grid test daily for 5 consecutives days. *Tibialis anterior* (TA) muscles were harvested approximately 2 hours after the last bout of exercise. **A**) qPCR analysis of *Fn14* mRNA expression in unexercised and grid test-exercised WT and *SOD1^G93A^* mice. Data are scatter dot plot mean L± SEM, *n* = 3-4 animals per experimental group, two-way ANOVA, ns = not significant, \*\**p* < 0.01. **B-H**) qPCR analysis of *Tweak* (**B**), *MuRF-1* (**C**), *Atrogin-1* (**D**), *Glut4* (**E**), Klf15 (**F**), *HKII* (**G**), *PGC-1*α (**H**) mRNA expression in unexercised and grid test-exercised WT, *Fn14^-/-^*, *SOD1^G93A^* and *SOD1^G93A^;Fn14^-/-^* mice. Data are scatter dot plot mean L± SEM, n = 4-9 animals per experimental group, two-way ANOVA, ns = not significant, \**p* < 0.05, \*\*\**p* < 0.001, \*\*\*\**p* < 0.0001.

Our results suggest that exercise regimens have a differential impact on the skeletal muscle of our 12-week-old experimental cohorts, pointing towards specific interactions between genotype, Fn14 depletion and exercise (Table 1).

**Table 1.** Effect of exercise and genotype on expression of the TWEAK/Fn14 signaling pathway and atrogenes.

|  | WT |  | *SOD1<sup>G93A</sup>* |  | *Fn14<sup>-/-</sup>* |  | *SOD1<sup>G93A</sup>;Fn14<sup>-/-</sup>* |  |
| --- | --- | --- | --- | --- | --- | --- | --- | --- |
|  | Rotarod | Grid test | Rotarod | Grid test | Rotarod | Grid test | Rotarod | Grid test |
| *Fn14* | Grey | Grey | Red | Blue | N/A | N/A | N/A | N/A |
| *Tweak* | Grey | Red | Grey | Grey | Grey | Grey | Red | Grey |
| *MuRF-1* | Grey | Grey | Red | Red | Grey | Grey | Grey | Grey |
| *Atrogin-1* | Grey | Grey | Red | Red | Red | Grey | Red | Grey |
| *Glut4* | Grey | Grey | Red | Grey | Grey | Grey | Grey | Grey |
| *Klf15* | Grey | Grey | Grey | Grey | Red | Grey | Grey | Red |
| *HKII* | Grey | Grey | Red | Grey | Red | Grey | Grey | Grey |
| *PGC-1α* | Grey | Grey | Red | Red | Grey | Grey | Red | Grey |

Red = upregulated
Blue = downregulated
Grey = no change

### Fn14 depletion abolishes the increase in myofiber size following exercise

We next determined if Fn14 depletion impacted myofiber size in the gastrocnemius muscle of rotarod- or grid test-trained 12-week-old WT, *Fn14^-/-^*, *SOD1^G93A^*and *SOD1^G93A^;Fn14^-/-^* males. Following rotarod training, we observed a significant increase in myofiber size of rotarod-trained WT animals compared to unexercised WT mice while this type of exercise did not impact myofiber size in *SOD1^G93A^* animals (Figure 7A). Interestingly, the myofiber size of rotarod-trained *Fn14^-/-^* and *SOD1^G93A^;Fn14^-/-^* mice was significantly smaller than in unexercised control animals (Figure 7A), indicating that the combination of Fn14 depletion and endurance exercise can reduce muscle size.

**Figure 7.**
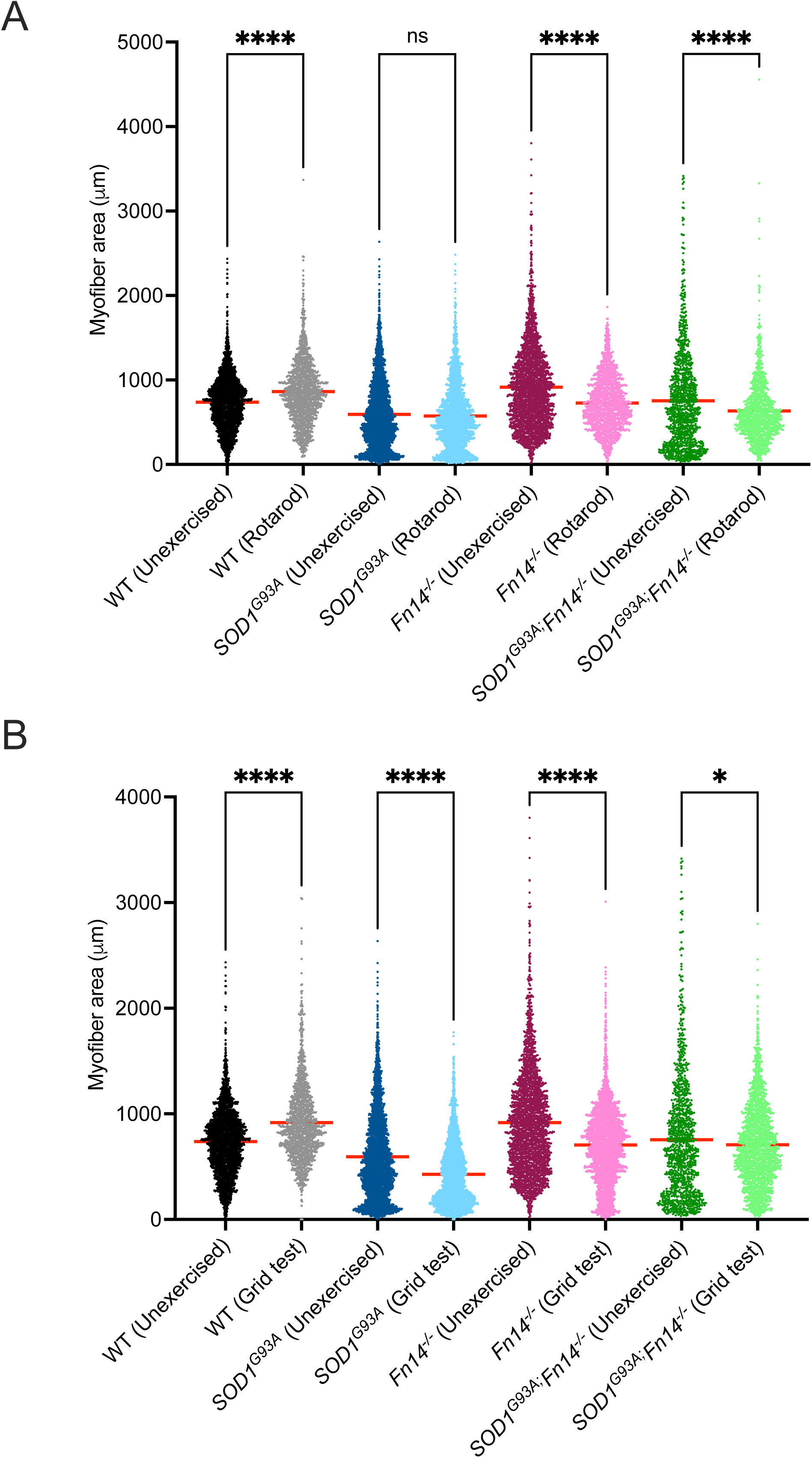
Combining exercise and Fn14 depletion has a negative impact on muscle fibre size. 12-week-old WT, *Fn14^-/-^*, *SOD1^G93A^* and *SOD1^G93A^;Fn14^-/-^*males either performed the grid test or were placed on the rotarod daily for 5 consecutives days. Gastrocnemius muscles were harvested approximately 2 hours after the last bout of exercise. **A**) Quantification of myofiber area of laminin-stained cross-sections of gastrocnemius muscles from 12-week-old unexercised and rotarod-exercised WT, *Fn14^-/-^*, *SOD1^G93A^*and *SOD1^G93A^;Fn14^-/-^* mice. Data are dot plot and mean, *n* = 3-4 animals per experimental group (>1100 myofibers per experimental group), two-way ANOVA, ns = not significant, \*\*\*\**p* < 0.0001. **B**) Quantification of myofiber area of laminin-stained cross-sections of gastrocnemius muscles from 12-week-old unexercised and grid test-exercised WT, *Fn14^-/-^*, *SOD1^G93A^* and *SOD1^G93A^;Fn14^-/-^* mice. Data are dot plot and mean, *n* = 3-4 animals per experimental group (>1100 myofibers per experimental group), two-way ANOVA, \**p* < 0.05.

Similar to rotarod-trained WT animals, grid test-trained WT mice displayed a significant increase in myofiber size compared to unexercised animals (Figure 7B). However, unlike rotarod-trained *SOD1^G93A^* mice, grid test-trained *SOD1^G93A^* animals had a significant decrease in myofiber size compared to unexercised *SOD1^G93A^* mice (Figure 7B). A significant decrease in myofiber area was also observed in grid test-trained *Fn14^-/-^*and *SOD1^G93A^;Fn14^-/-^* mice compared to unexercised control animals (Figure 7B), suggesting that both the *SOD1^G93A^* genotype and Fn14 depletion negatively impact muscle size following a resistance exercise regimen.

Together, our data points to independent influence of exercise type and genetics in muscle fiber size.

## DISCUSSION

In this study, we aimed to better understand how increased Fn14 expression in an ALS mouse model with chronic denervation and muscle wasting could contribute to muscle pathology and disease progression. To achieve this, we genetically deleted *Fn14* in both WT and *SOD1^G93A^* mice and observed behavioural, molecular and histological changes that were dependent on exercise and disease progression.

In the first instance, we not only confirmed our previous observation of increased expression of Fn14 in the skeletal muscle of *SOD1^G93A^*ALS mice during disease progression [33], but we also validated the previously reported negative correlation between the activity of the TWEAK/Fn14 pathway and the expression of the metabolic effectors *Glut4*, *Klf15*, *HKII* and *PGC-1*α [29]. Interestingly, we recently demonstrated a similar but inverse negative correlation in the skeletal muscle of another neuromuscular mouse model, SMA mice, whereby the expression levels of *Tweak* and *Fn14* decreased during disease progression while those of *Glut4*, *Klf15*, *HKII* and *PGC-1*α increased [32]. One important distinction between these two studies is the developmental stage investigated. Indeed, SMA mice were of pre-weaned age [32] while the *SOD1^G93A^* ALS mice were at adult stages (current study), suggesting that the TWEAK/Fn14 signalling pathway is differentially regulated at different stages of muscle development. This differential regulation might have an impact on downstream metabolic requirements and regulation as well as therapeutic interventions in cases of dysregulation.

One of our key findings is the extended lifespan of Fn14-depleted *SOD1^G93A^* ALS mice, which is contrary to the absence of impact following genetic *Tweak* deletion in the same mouse model, which we have previously reported [33]. This suggest that the detrimental effect of the aberrant activity of the TWEAK/Fn14 pathway in skeletal muscle of *SOD1^G93A^* ALS mice is driven by the receptor (Fn14) and not the ligand (TWEAK). This aligns with previous work that points to a greater role for Fn14 than TWEAK in enabling pathway activity [34]. It is also possible that the differential impacts observed in TWEAK- and Fn14-depleted *SOD1^G93A^* ALS mice are due to Fn14-independent Tweak signalling [50] and/or TWEAK-independent Fn14 signalling [51]. Furthermore, the distinct effects of TWEAK and Fn14 depletion in *SOD1^G93A^* ALS mice could further be caused by their known roles in other tissues such as the heart, gastrointestinal tract, kidney, liver, central nervous system and epithelium [52–54]. As the genetic knock-out of Tweak and Fn14 was systemic in both cases, we cannot exclude additional benefits or detrimental effects stemming from altered function in other cells and tissues. Regardless of the reasons, our combined studies point to a greater therapeutic value in modulating Fn14 over TWEAK.

In addition to lifespan, we also observed improvements in skeletal muscle pathology at molecular and histological levels in Fn14-depleted *SOD1^G93A^* mice. These changes did not occur in *Fn14^-/-^* mice when compared to WT animals, suggesting that the effects were dependent on disease stage. Interestingly, we have previously shown increased muscle fibre and NMJ endplate sizes in TWEAK-depleted *SOD1^G93A^* ALS mice [33], further supporting a role for the TWEAK/Fn14 pathway in muscle pathology in this mouse model and in more general adult denervation-induced muscle atrophy [34]. Of note is that in both TWEAK- and Fn14-depleted *SOD1^G93A^* ALS mice, there were no significant improvements in motor function [33], suggesting that simply targeting the TWEAK/Fn14 pathway is not sufficient for the recovery of the neuromuscular unit.

Surprisingly, the beneficial impact of Fn14 depletion on the survival of *SOD1^G93A^* ALS mice was almost masked when the mice underwent weekly rotarod and grid test assessments for approximately 16 weeks as the enhanced physical activity itself had a positive impact on survival of the *SOD1^G93A^* ALS mice. Further investigations showed that the combination of 5 consecutive days of exercise (rotarod or grid test) and Fn14 depletion was sufficient to induce changes at molecular and histological levels in the skeletal muscle of 12-week-old animals. These changes were dependent on both disease stage and exercise, with the rotarod representing an endurance exercise and the grid test a resistance exercise. Combining exercise and Fn14 depletion may therefore lead to potentially, additive, synergistic and/or antagonistic interactions that may be dependent on the exercise regimen itself and state of the disease.

One key observation was that changes in *Tweak* and *Fn14* expression appeared dependent on the type of exercise and genotype of the animal. *Fn14* levels displayed a differential expression in *SOD1^G93A^* mice only, whereby it was increased following rotarod and decreased following the grid test. *Tweak* expression however was only increased in *SOD1^G93A^;Fn14^-/-^*mice after the rotarod and in WT animals after the grid test, when compared to unexercised controls. These diverse patterns may reflect the complex metabolic adaptations impacted by disease, Fn14 presence/absence and type of exercise. Typically, endurance exercises promote the use of aerobic/oxidative metabolic pathways in skeletal muscle while resistance exercises favour anaerobic/glycolytic metabolic pathways [55]. In ALS, skeletal muscle metabolism during rest and exercise is altered in both pre-clinical models and patients [56–59], which could alter how ALS muscle adapts to different types of exercises and the overall beneficial vs detrimental outcomes [60]. As for Fn14, it typically increased in skeletal muscle of healthy individuals and adult mice following exercise, irrespective of type (endurance vs resistance) [61–64]. Conversely, the muscle-specific deletion of Fn14 and the ubiquitous TWEAK deletion in mice both improved exercise capacity and oxidative metabolism [65,66], suggesting that sustained and/or aberrant increase in TWEAK/Fn14 activity expression during exercise may be detrimental. It is therefore unclear why the expression of both the ligand and effector are commonly reported as being elevated following exercise. Of note, we did not observe changes in *Fn14* expression in exercised WT mice in our study, which may be due to our selected exercise regimens (length and type of exercise). Nevertheless, our results, combined with previous studies, suggest and support a complex interaction between Fn14 regulation, disease state, exercise and the metabolic status of muscle.

Another noticeable result is the influence of genotype and exercise on the expression of the atrogenes *MuRF-1* and *Atrogin-1*. Indeed, we found that the expression of *MuRF-1* is significantly elevated in *SOD1^G93A^* mice following both the rotarod and grid test, supporting previous studies on the negative impact of exercise in ALS patients [67,68]. In *SOD1^G93A^;Fn14^-/-^*mice however, *MuRF-1* levels remained low in both rotarod and grid test groups, aligning with the previous report of reduced neurogenic muscle atrophy in muscle-specific Fn14-depleted animals [65]. Interestingly, *Atrogin-1* levels were also significantly increased in *SOD1^G93A^* mice after the rotarod and grid test. However, *Atrogin-1* levels were also upregulated in *Fn14^-/-^*mice but only after the rotarod and not after the grid test. Furthermore, *Atrogin-1* expression remained elevated in *SOD1^G93A^;Fn14^-/-^*mice after the rotarod while remaining low after the grid test. While the differential expression patterns of both atrogenes might appear contradictory, previous studies have demonstrated that their regulation can be controlled by distinct pathways [69,70]. Of note, our analysis of muscle fibres shows that while both exercise regimens led to a significant increase in myofiber size in WT mice, there was an overall decrease in the other three experimental groups. This suggests that changes in *MuRF-1* and *Atrogin-1* are not sufficient to improve muscle size and that other molecular effectors and regulatory pathways may be responsible for modulating muscle mass [71].

Finally, the expression of metabolic effectors previously shown to be regulated by TWEAK/Fn14 signalling also appear to be dependent on genotype and type of exercise. For example, *Glut4* expression is specifically increased in rotarod-exercised *SOD1^G93A^* mice only, suggesting that it is induced by endurance exercises and somewhat modulated by Fn14 as *Glut4* levels remain low in rotarod exercised *SOD1^G93A^;Fn14^-/-^*mice. Another example is *PGC-1α*, where both the rotarod and grid test result in its upregulation in *SOD1^G93A^* mice and Fn14 depletion restores the levels to normal only in the grid-test exercised *SOD1^G93A^;Fn14^-/-^*mice. These differential patterns and relationships between genotype, exercise and metabolic effectors most likely result from the combination of different metabolic pathways favoured by different types of exercise [55] and the impact of the ALS-causing mutations on muscle metabolism [56–59].

While our work provides some interesting insights, it is important to note its key limitations. Firstly, the impact of exercise on *SOD1^G93A^;Fn14^-/-^* mice was observed in animals that performed both types of exercise weekly from 8 weeks of age to humane endpoint. However, the rotarod and grid test experiments were done separately on 12-week-old animals for 1 week only. Furthermore, our study focused on the known metabolic effectors downstream of TWEAK and Fn14, which means that additional genes and signalling cascades could be impacted by exercise and/or genotype and contribute to our observed results. Finally, our research was aimed at investigating skeletal muscle but as Fn14 depletion is systemic, some of the beneficial and detrimental effects reported may be due to other cells and tissues.

## CONCLUSIONS

Our study provides additional insights on the role of the TWEAK/Fn14 pathway in a denervation-induced muscle pathology as modelled in the *SOD1^G93A^* ALS mice. Importantly, we demonstrate that the benefits of Fn14 depletion are impacted by exercise. This is particularly relevant in the context of the current therapeutic landscape of the ALS field, where combinatorial therapies that include exercise regimens are being explored by many research and clinical teams. As such, a better understanding and consideration of the interactions between treatments, muscle metabolism and exercise will be of importance in future studies.

## LIST OF ABBREVIATIONS

α-BTX: α-bungarotoxin
ALS: Amyotrophic lateral sclerosis
C9ORF72: chromosome 9 open reading frame 72
FBS: fetal bovine serum
Fn14: fibroblast growth factor inducible 14
FUS: Fused in Sarcoma
Glut4: glucose transporter 4
HKII: hexokinase II
Klf15: Krüppel-like factor 15
NF-κB: nuclear factor kappa-light-chain-enhancer of activated B cells
PBS: phosphate buffered saline
PFA: paraformaldehyde
PGC-1α: peroxisome proliferator-activated receptor-gamma coactivator 1α
PolJ: RNA polymerase II polypeptide J
SDS: sodium dodecyl-sulfate
SMA: spinal muscular atrophy
SOD1: superoxide dismutase 1
TA: tibialis anterior
TDP-43: TAR DNA-binding protein 43
TWEAK: tumor necrosis factor-like weak inducer of apoptosis

## DECLARATIONS

### Ethics approval and consent to participate

Not applicable.

### Consent for publication

Not applicable

### Availability of data and materials

All data generated or analysed during this study are either included in this published article [and its supplementary information files] or are available from the corresponding author on reasonable request.

### Authors’ contributions

Conceptualisation: G.H., C.R., M.B.; Methodology : G.H., M.B.; Validation : G.H., M.B.; Formal analysis: G.H., N.A., E.R.S., M.O., J.M.H., T.S.,.S.I., A.B., A.K., F.S., M.B.; Investigation: G.H., N.A., E.R.S., M.O., J.M.H., T.S.,.S.I., E.M., A.B., A.K., F.S., M.B.; Writing-original draft preparation: M.B.; Writing-review and editing: G.H., N.A., E.R.S., M.O., J.M.H., T.S.,.S.I., E.M., A.B., A.K., F.S., C.R., M.J.A.W., M.B.; Visualisation: M.B.; Supervision: C.R., M.J.A.W., M.B.; Project administration: G.H., M.B.; Funding acquisition: C.R., M.J.A.W., M.B.

## Acknowledgements

We would like to thank the staff at the BMS facility at the University of Oxford, Dr Linda Burkly and Biogen.

## SUPPLEMENTARY TABLES

**Supplementary Table 1.** Mouse primers used for quantitative real-time PCR.

