## Supplementary Table 1 for "Exercise and disease state influence the beneficial effects of Fn14-depletion on survival and muscle pathology in the *SOD1^G93A^* amyotrophic lateral sclerosis (ALS) mouse model"

Supplementary Table 1. Mouse primers used for quantitative real-time PCR.

| Gene | Forward | Reverse |
| --- | --- | --- |
| *Atrogin-1* | 5’-TCAAAGGCCTCACGATCACC-3’ | 5’-CCTCAATGACGTATCCCCCG-3’ |
| *Fn14* | 5′-TCGTGTTGGGATTCGGCTTGGT-3' | 5′-ACTTTTCTCTCCGGCGGCATCT-3' |
| *Glut4* | 5′-GACGGACACTCCATCTGTTG-3' | 5′-CATAGCTCATGGCTGGAACC-3' |
| *HKII* | 5′-GAAGGGGCTAGGAGCTACCA-3' | 5′-CTCGGAGCACACGGAAGTT-3' |
| *Klf15* | 5′-TGCGTCGGCACACAGGCGAGAA-3' | 5′-CCGGTGCCTTGACAACTCATCT-3' |
| *MuRF-1* | 5′-AGGACTCCTGCCGAGTGAC-3' | 5′-TTGTGGCTCAGTTCCTCCTT-3' |
| *PGC-1α* | 5′-TGGAGTGACATAGAGTGTGCTGC-3' | 5′-CTCAAATATGTTCGCAGGCTCA-3' |
| *PolJ* | 5′-ACCACACTCTGGGGAACATC-3' | 5′-CTCGCTGATGAGGTCTGTGA-3' |
| *Tweak* | 5′-AAGTTCACTGAGGGGCCTTGCT-3' | 5′-TGTGAACAAGCTCTGGCTGCCT-3' |
